# Social Perception of Young Adults Prolongs the Lifespan of Aged Drosophila

**DOI:** 10.1101/2020.08.30.273854

**Authors:** Li-Chun Cho, Chih-Chieh Yu, Chih-Fei Kao

**Affiliations:** Department of Biological Science and Technology, College of Biological Science and Technology, National Chiao Tung University, Hsinchu, Taiwan; Center for Intelligent Drug Systems and Smart Bio-devices (IDS2B), National Chiao Tung University, Hsinchu, Taiwan; Institute of Molecular Medicine and Biochemical Engineering, National Chiao Tung University, Hsinchu, Taiwan

**Keywords:** social perception, social surroundings, chemosensory cues, olfaction, gustation, lifespan modulation, Drosophila melanogaster

## Abstract

Lifespan is modulated at distinct levels by multiple factors, including genetic backgrounds, the environment, behavior traits, the metabolic status, and more interestingly, the sensory perceptions. However, the effects of social perception between individuals living in the same space remain less clear. Here, we used the Drosophila model to study the influences of social perception on the lifespan of aged fruit flies. We found the mean lifespan of aged Drosophila is markedly prolonged after being co-housed with young adults of the same gender. Moreover, the changes of lifespan were affected by several experimental contexts: (1) the ratios of aged and young adults co-housed, (2) the chronological ages of two populations, and (3) the integrity of sensory modalities. Together, we hypothesize the chemical/physical stimuli derived from the interacting young adults are capable of interfering with the physiology and behavior of aged flies, ultimately leading to the alteration of lifespan.

## Introduction

Lifespan is modulated by diverse factors that mostly act through the alteration of internal physiology and the initiation of behaviors that respond to the external environment (1-4). Moreover, the capability to modulate organism lifespan by various genetic and non-genetic manipulations further indicate the plasticity and limitations that apply on the life expectancy of organisms. Intriguingly, recent studies have demonstrated manipulations in the sensory systems, which provide organisms with the ability to perceive and interact with the internal and external environments, are capable of altering organism lifespan, highlighting the potential linkages between sensory perception and aging processes (5, 6). Such an example includes the loss of olfaction, which is the sense of smell that allows organisms to perceive the chemical landscapes of external environment, significantly influences the lifespan of model organisms, such as worms and fruit flies (7, 8). Extended studies further indicate the plausible connections between the olfactory sensation, the metabolic changes, and ultimately the longevity of organisms (9). Likewise, increasing evidence suggests the activity of additional sensory modalities also have the capability to differentially influence the physiology and the longevity in animals across taxa (for reviews, see 5, 6). In the real world, the perception of both external and internal states usually involves multiple sensory-related cells/molecules. Among the distinct perceptions, social perception apprehends social cues emanated from the conspecifics living in the same space and recognizes the changes of social surroundings. For the past years, the influences of social surroundings have on overall health of organisms are well noted from the human-based clinical and epidemiological studies. Both types of studies suggest favorable social experiences are associated with the increase of well-being and healthy aging (6, 10). However, to better understand the physiological impacts of social perception in different social contexts and the underlying neural networks/molecules, small model organisms, such as fruit flies, kept in distinct social situations provide a convenient experimental platform.

Here, we used the model organism, Drosophila melanogaster, to explore the physiological impacts to the aged fruit flies when experiencing distinct contexts of social surroundings. More specifically, we focused on the effects conferred by the co-housed young fruit flies. In addition, we further elucidated the sensory pathways in the aged flies to fully perceive the social cues from the co-housed young flies.

## Results

### The life expectancy of aged flies is differentially affected by the social surroundings

To explore whether the presence of young individuals in the same living environment could affect the lifespan of aged individuals, we used Drosophila melanogaster as a model system to create such scenery. Unmated wild type (WT) flies were cultured according to the standard conditions (see Methods) and were housed at a fixed population size (20 flies/vial). Given the lifespan of fruit flies is profoundly affected by the sex experience/behavior (11-14), our co-housing experiments only include flies of the same gender to avoid such influences (Fig. 1A). At the age of 40 days (slightly earlier than the median lifespan of Canton-S flies; Fig. S1A), marked aged flies were relocated to live with 1 day (1d)-old unmated young adults for the rest of their lives. As shown in Fig. 1B, the mean lifespan of aged Canton-S flies was notably affected by co-housing with 1d-old young adults. In our pilot experiments, we kept the ratio of aged vs. young flies at 1:3 to analyze the effects of young adults to aged flies. Intriguingly, in general, the mean lifespan of aged flies showed 14∼25% extension compared to controls (Fig. 1B; ratio 1:3, aged vs. young; please note that, in the control experiments, the population size of 40d-old flies was kept at 20 flies/vial, same as the experimental groups, to minimize the influences of population density). Similar lifespan extension phenotypes were also observed in two additional WT Drosophila strains (Figs. S1B-S1D; Oregon-R and w^1118^; the effects were not significant in Oregon-R male flies). Moreover, the mean lifespan of young flies co-housed was not affected by the presence of aged animals in all three WT fly strains we tested (Fig. S2). Increasing the density of young flies in the co-housing experiments further boosted the lifespan extension phenotypes in male flies, but not in female flies (Fig. 1B; ratio 1:9, aged vs. young). Overall, regardless the gender differences, the mean lifespan of aged fruit flies was prolonged when residing with an exceeding number of young adults. Next, we were interested to know whether the lifespan extension phenotypes persist when equal amount of aged and young flies were present in the same living environment (Fig. 1B; ratio 1:1, aged vs. young). Under this condition, the mean lifespan of experimental aged flies was comparable to WT flies in both genders. Moreover, in the condition that the density of co-cultured young adult Drosophila was decreasing to a 3:1 ratio (aged vs. young), the mean lifespan of aged female flies was slightly reduced, while aged males were less affected (Fig. 1B). Together, our results suggest aged flies recognize and respond to the changes of social surroundings, which may directly or indirectly derive from the co-housed unmated 1d-old young fruit flies. We therefore called this longevity promoting effects, the “youth impacts” (Fig. 1C). In line with our findings, if the 40d-old aged flies could continue to be surrounded by the fresh batches of 1d-old young flies, was it possible to have stronger longevity promoting effects (youth impacts) to aged flies? This scenario was achieved by replacing the batch of co-housed 1d-old fruit flies every 7 days (Fig. 1D). Surprisingly, even though the lifespan extension phenotypes remained, the overall effectiveness, regardless of genders, was not as good as the condition that only one batch of young flies was co-housed with the aged flies (Fig. 1E). The inability of fresh youth impacts to further promote the longevity of aged flies suggests a probable limit of longevity extending capability. Another conceivable explanation highlights the importance of accumulating sufficient strength of social perception to the co-housed young adults. It is likely that aged flies require more than 7 days to make beneficial social connections with young flies living in the same space.

**Fig. 1.**
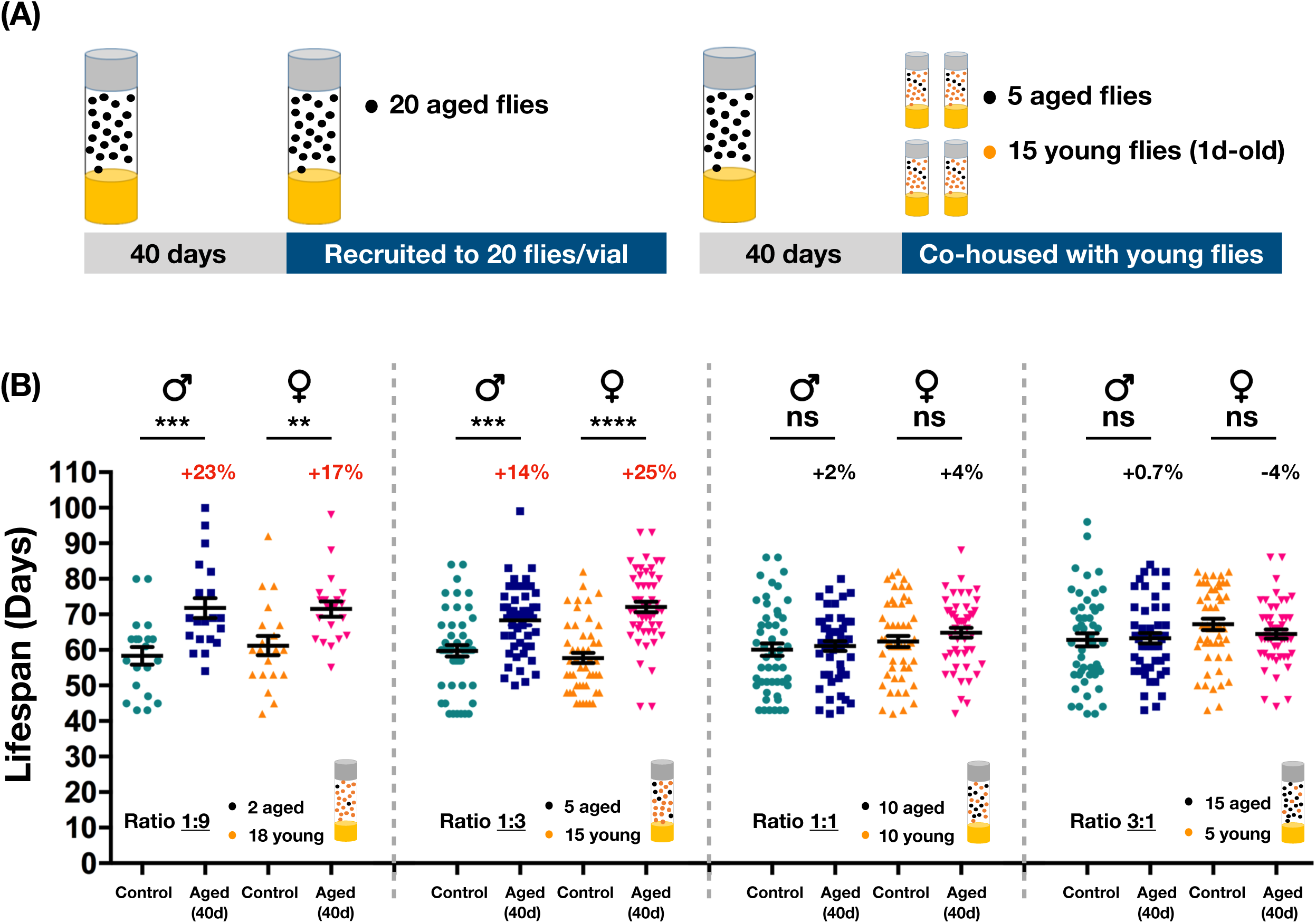
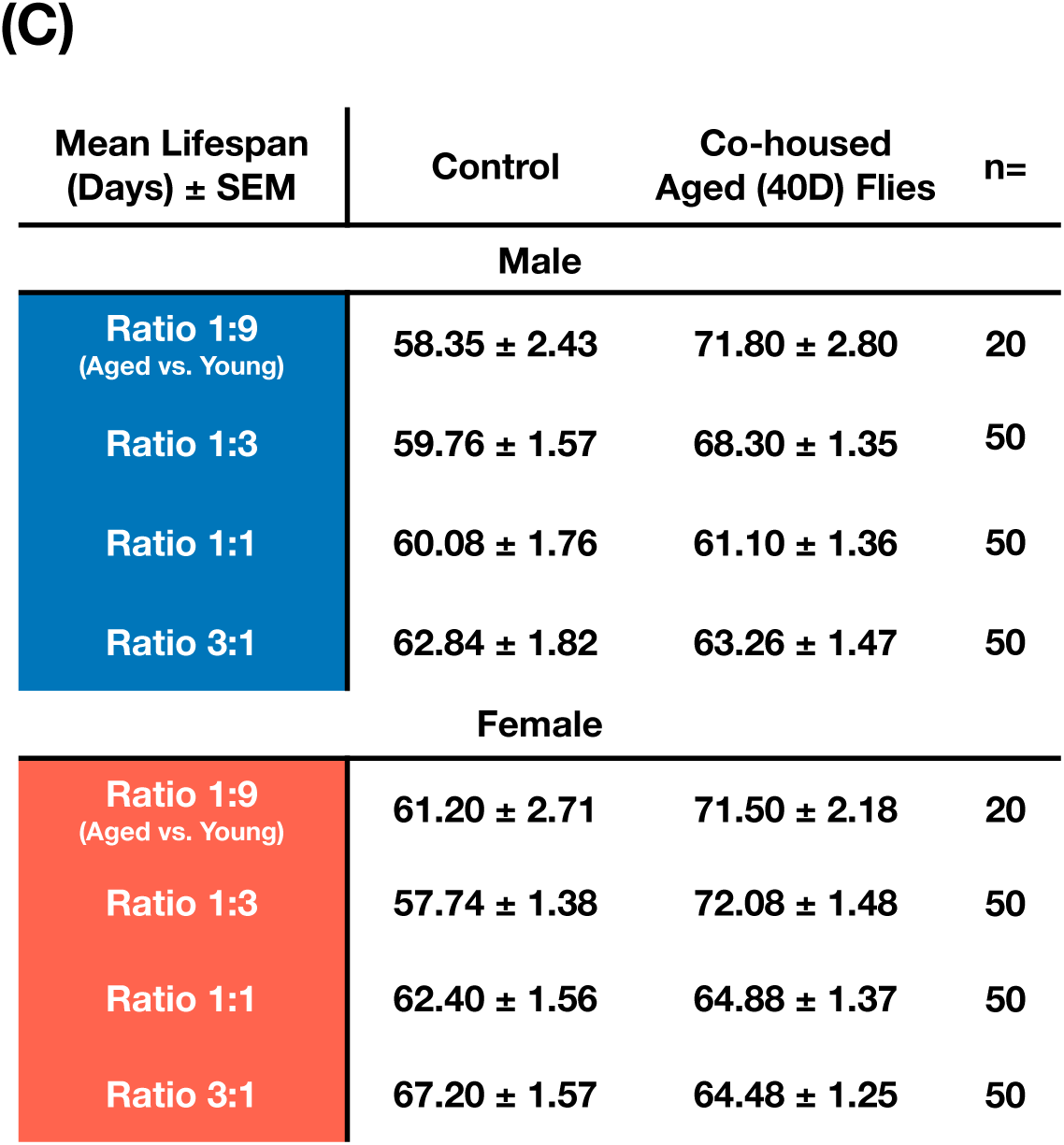
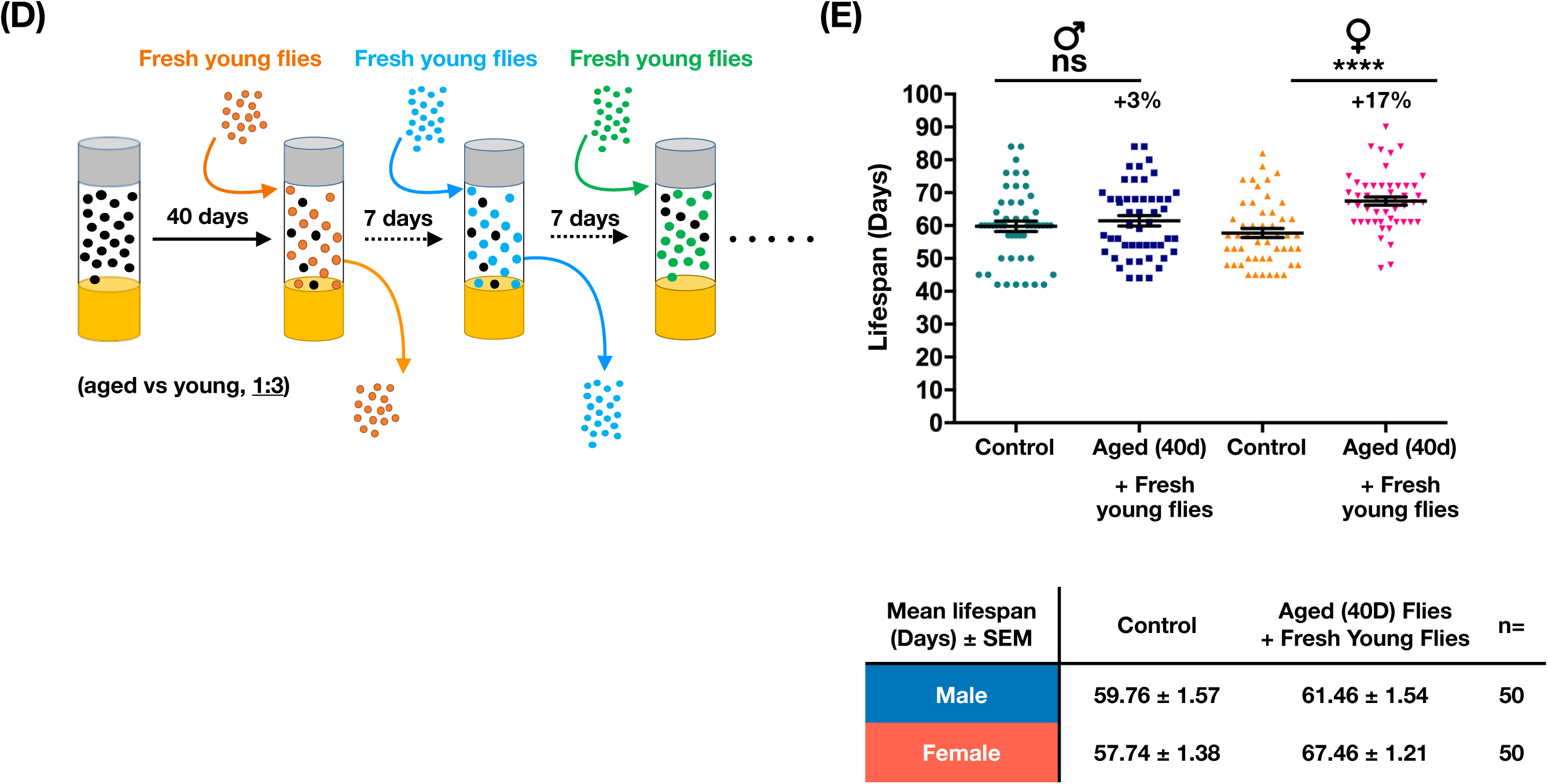
The lifespan of aged flies is altered by co-housing with young adults. **(A)** An illustration of the co-housing experiments. To keep the population density constant, in the control experiments, the number of 40d-old aged flies was maintained at 20 flies/vial. **(B)** Canton-S aged (40d-old, unmated) flies were co-housed with young adult flies (1d-old, unmated; same gender) in different cohort ratios (aged:young = 1:9, 1:3, 1:1, 3:1). When co-housed with an exceeding number of young flies, aged flies showed significant lifespan extension (aged:young ratio 1:9, n=20 for each column; 1:3, n=50 for each column). The mean lifespan of aged flies had no significant changes after being co-housed with an equal or a less number of young flies (aged:young ratio 1:1 and 3:1; each column contains the results of 50 flies). Results were expressed as means ± SEM. **p<0.01, ***p<0.001, ****p<0.0001, according to the Mann-Whitney test. Percentage changes of mean lifespan were also indicated. **(C)** Summary of the lifespan results shown in (B) and Fig. S1B and S1C. **(D)** The experimental scheme of co-housing assays, in which the same five 40d-old Canton-S aged flies was co-housed with a fresh batch of 1d-old (15 flies) every 7 days, was illustrated. Lifespan results were expressed as means ± SEM and shown in **(E)**. **** p<0.0001, p value is determined by the Mann-Whitney test. Each column contains the results of 50 flies. Percentage changes of mean lifespan were indicated at the top.

Next, to study if the youth impacts were also effective on aged flies that are slightly younger/older than 40 days, similar co-housing experiments were performed accordingly. While similar lifespan extension phenotypes were observed, the overall effects on the 40d-old adult Drosophila were more profound in comparison to 30d- and 50d-old flies (Fig. 2). These results suggest there may be a best chronological window for the aging flies to receive the youth impacts from co-housed young adults. However, it is currently not clear why 40d-old fruit flies have the utmost responses to the youth impacts. The following interesting question we asked was what chronological ages of young flies are able to confer the pro-longevity signals. To this end, 40d-old flies were arranged to live with the same gender, unmated flies of selected chronological ages younger than 40 days (Fig. 3A; 1d-, 10d-, 20d-, and 30d-old flies). Our results showed that aged flies perceive the most notable lifespan promoting signals from the youngest flies (i.e., co-housing of 40d-old and 1d-old flies; the age difference is 39 days; Figs. 3B-3D). Strikingly, the lifespan extension phenotypes were almost completely lost in 40d-old male flies that were co-housed with 10d-old unmated males, as well as 20d- and 30d-old adults (Fig. 3B). However, unlike the male flies, aged female flies appeared to be less sensitive to the chronological ages of co-housed younger females. In all conditions we tested, the mean lifespans of aged female flies were readily prolonged, albeit the effects were less potent (Fig. 3C). In summary, our results suggest the deterioration of youth impacts occurs quickly within less than 10 days (Fig. 3D).

**Fig. 2.**
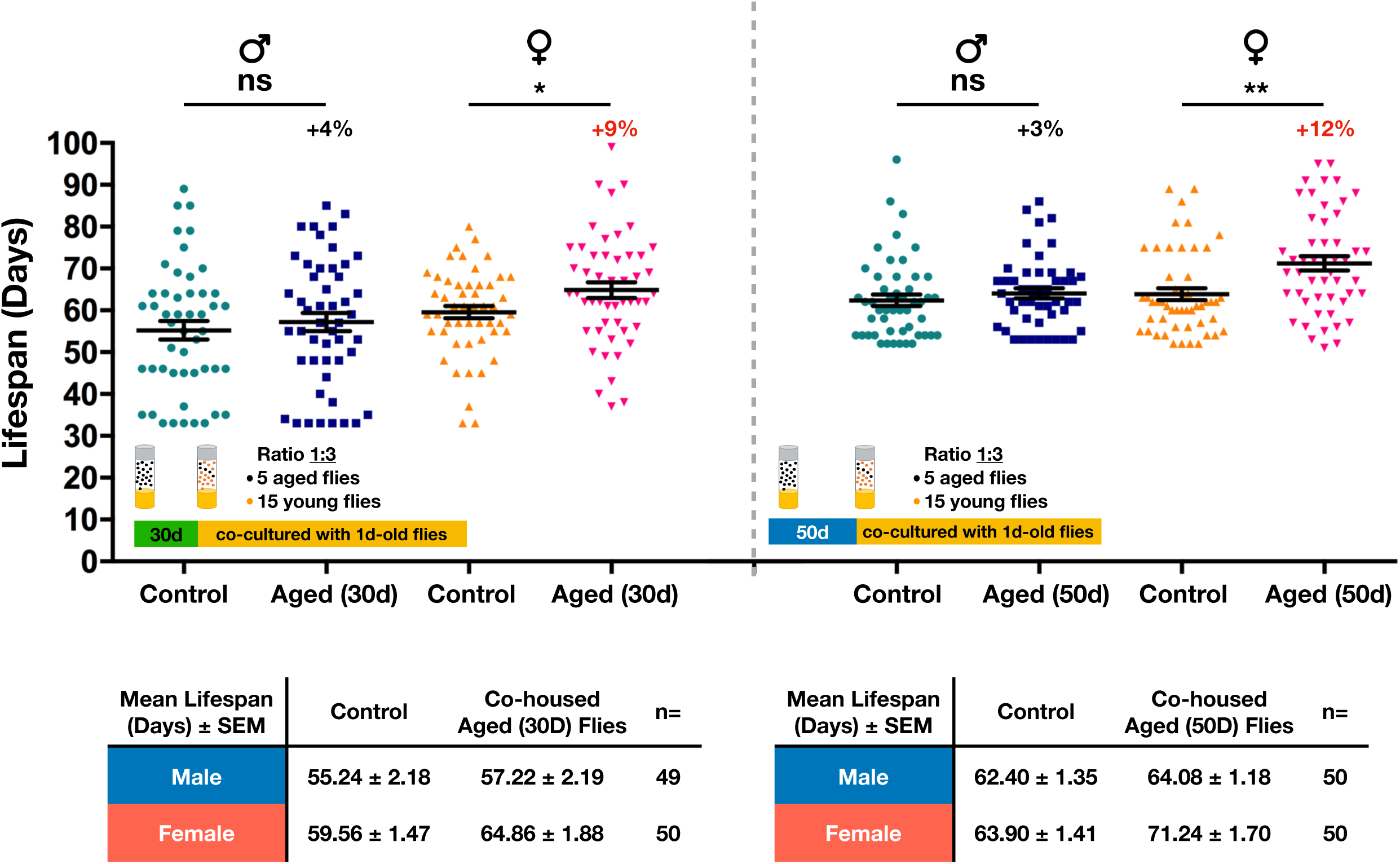
Lifespan extension phenotypes are affected by the chronological age of aged flies. Canton-S flies of indicated chronological ages (30d- and 50d-old) were co-housed with 1d-old flies at the cohort ratio of 1:3. Each column contains the results of 49∼50 flies. Results were expressed as means ± SEM and summarized at the bottom. In call cases, the longevity promoting effects were more robust in female flies. *p<0.05, **p<0.01, according to the Mann-Whitney test. ns: not significant. Percentage changes of mean lifespan were indicated on the top.

**Fig. 3.**
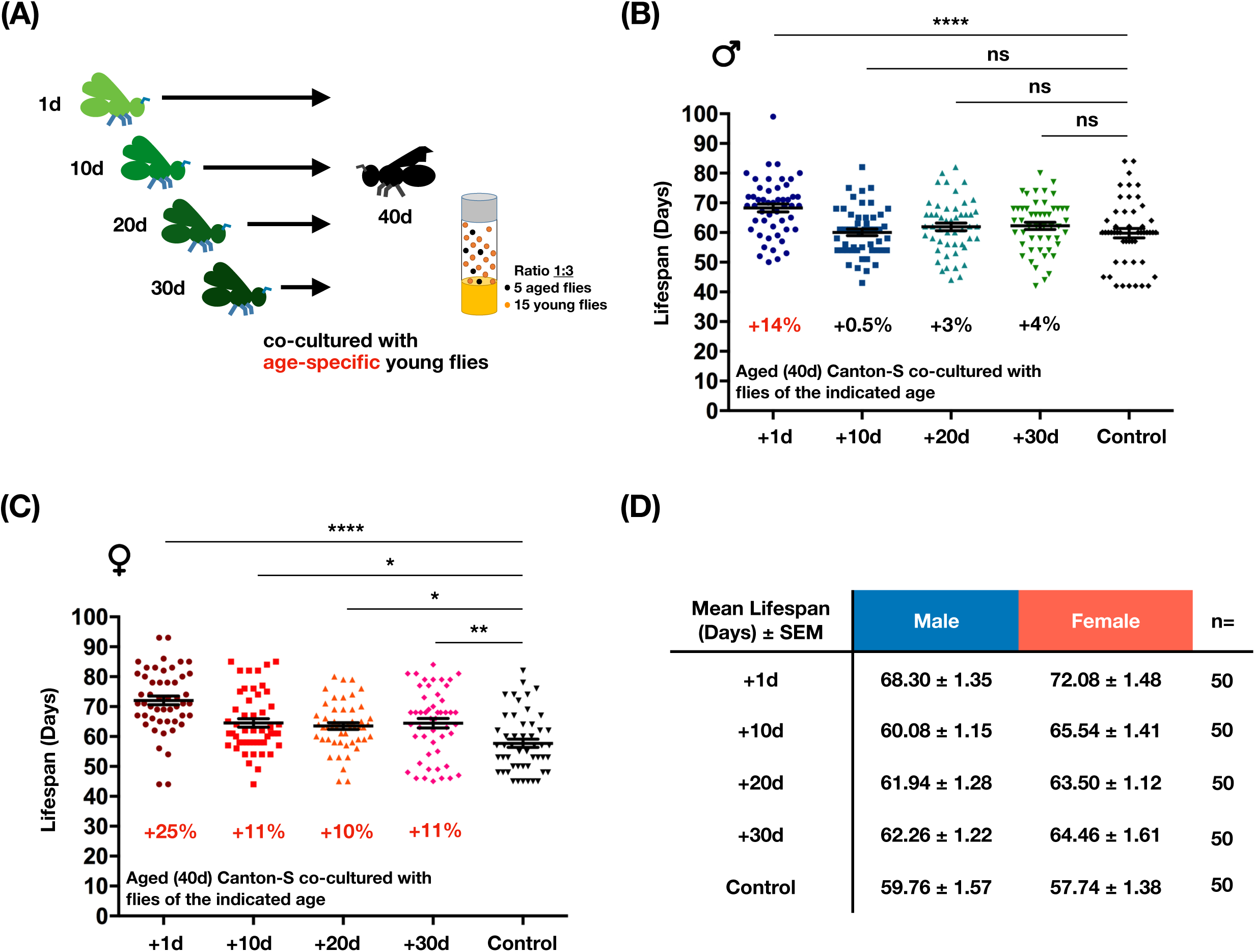
Chronological ages of the co-housed young flies have different longevity promoting capabilities. **(A)** An illustration of the experimental concept, in which aged flies were co-cultured with age-specific young flies, was delineated. Briefly, 40d-old Canton-S aged flies were co-housed with 1d-, 10d-, 20d-, 30d-old young flies respectively. Results of lifespan were shown in **(B)** for male flies, and **(C)** for female flies. In all conditions, aged flies acquired stronger lifespan extension phenotypes when co-housed with younger flies. *p<0.05, **p<0.01, ****p<0.0001, group compared by the Kruskal-Wallis test and followed by Dunn’s multiple comparison test (each column contains 50 flies). Percentage changes of mean lifespan were indicated at the bottom. ns: not significant. **(D)** Summary of results shown in (B) and (C).

### Social perception involves the olfactory and gustatory systems

In considering the olfactory sensation, which mediates the detection and recognition of volatile compounds, is one of the most important sensations in insects (15-17), we were interested to know whether olfaction is involved in perceiving the youth impacts. Flies with defective *or83b* gene are anosmic and live longer, especially in the mutant female flies (Fig. 4A) (8, 18, 19). Given the longer lifespan of *or83b* mutant flies, we included the 60d-old, but not the 40d-old, mutants in the co-housing experiments. As shown in Fig. 4B, the mean lifespan of aged *or83b* mutants was not significantly extended after being co-housed with an exceeding number of 1d-old WT adults. Our results therefore suggest olfaction may be involved in the perception of youth impacts. However, in addition to the olfaction, the gustation also provides another spectrum of environmental cues to the organisms, such as pheromones and trace of foods (20-22). Different from the mechanisms of olfaction, it is experimentally challenging to eradicate gustatory perception entirely due to the presence of diverse gustation-related receptors and lack of common regulatory mechanism (23). Physical ablation of labellum, legs, and wings, which house most of the gustatory neurons, dramatically weakened the flies, leading to organism death within a few days. We therefore selectively tested the *ppk23* (pickpocket 23) mutants in the co-housing experiments. Ppk23, a member of the Degenerin/Epithelial sodium channel (DEG/ENaC) family, is a gustatory-specific putative non-voltage gated cation channel and is broadly expressed in taste-related neurons (24-27). The survival assays indicated the lifespan of *ppk23*^*-/-*^ female flies, but not male flies, was longer than w^1118^ controls (Fig. 4A). Therefore, similar to the *or83b* mutants, 60d-old *ppk23*^*-/-*^ flies were assayed in the co-housing experiments. As shown Fig. 4C, we surprisingly found the mean lifespan of aged *ppk23* (60d-old) mutants was also not significantly extended after being co-housed with an exceeding number of 1d-old w^1118^ adults. Together, our results therefore suggest the functional integrity of both olfaction and gustation may be critical for aged flies to perceive and recognize the youth impacts. Loss of either sensory modality led to the impaired ability to fully appreciate the changes of social surroundings. To explore the identity of the youth impacts, we collected the cuticular extracts (including cuticular hydrocarbons, pheromones, and other organic chemicals that were dissolved by the hexane solution) from 1∼2d-old young flies, since social interaction between flies mostly involves chemical communications (27-30). To rid of other external sources of cuticular chemicals, only a single fly was cultured in a space-restricted vial containing a piece of cuticular extract-rinsed filter paper (Fig. S3A). Reduction of living space was to increase the frequency of contact to the cuticular extracts. Moreover, to maintain the efficacy of cuticular extracts, the fly was transferred to a new vial with fresh cuticular extracts every two days. As suspected, the exposure to hexane alone reduced the lifespan of adult flies (+Hexane) when comparing to mock controls (Control; Fig. S3B). However, flies living in the cuticular extract-conditioned vial had slightly longer mean lifespan than the hexane-only group. Overall, the pro-longevity effects were more robust in male flies, suggesting the strength of youth impacts may partly derive from chemical signals transferred from the co-housed young flies. However, the low efficacy of cuticular extracts may result from the low frequency of physical contacts with the filter paper or the insufficient amount of cuticular extracts applied. Moreover, the cuticular extracts isolated by the organic solvent may not represent all the chemicals that could be found on the body of fruit flies, since lots of the cuticular chemicals are volatile and have relatively short half-life. In addition to the chemical communications, physical engagement with the young flies may also partly contribute to the effects of youth impacts.

**Fig. 4.**
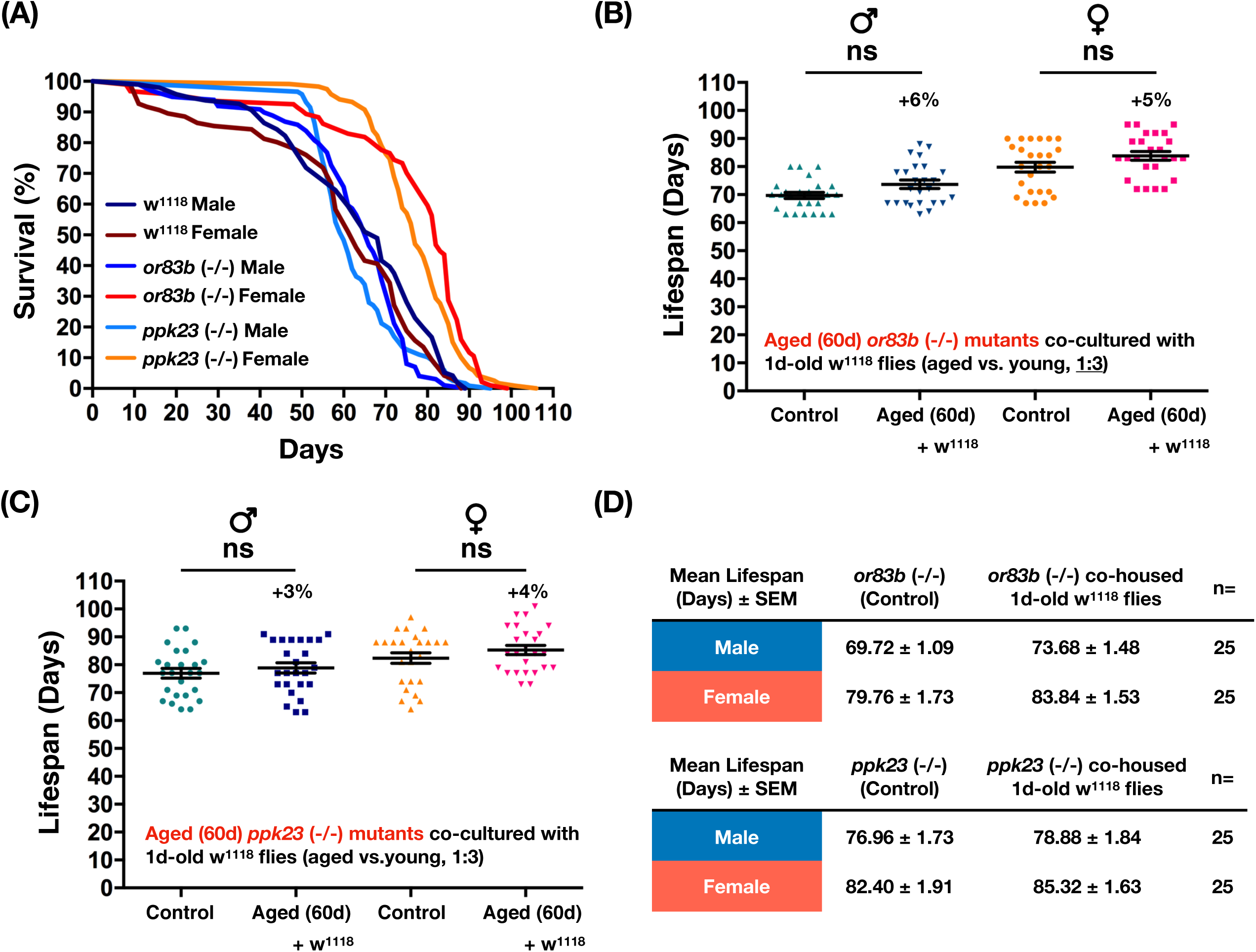
Social perception of co-housed young flies by the aged flies involves the sensory inputs from olfaction and gustation. **(A)** Survival curves of the *or83b(-/-), ppk23(-/-)*, and w^1118^ control flies were shown. All flies were maintained at the density of 20 flies per vial. Median lifespan for each genotype is: (1) w^1118^ male: 65 days; n=100, (2) *or83b(-/-)* male: 66 days; n=100, (3) *ppk23(-/-)* male: 60 days; n=118, (4) w^1118^ female: 61 days; n=100, (5) *or83b(-/-)* female: 82 days; n=96, (6) *ppk23(-/-)* female: 77 days; n=119. **(B)** and **(C)**, 60d-old aged *or83b(-/-)* flies or *ppk23(-/-)* flies were co-housed with 1d-old w^1118^ flies in a 1:3 ratio. Lifespan of individual flies was recorded and analyzed. p values were annotated by the Mann-Whitney test. Each column contains results from 25 flies. Percentage changes of mean lifespan were indicated on the top. In all cases, p values were greater than 0.05. ns: not significant. **(D)** Summary of results shown in (B) and (C).

## Discussion

Besides the pro-longevity signals, the youth impacts may provide additional beneficial effects to various disease conditions. For example, a previous study has demonstrated social interactions with co-housed young flies are able to improve the physiological defects, such as the reduction in lifespan, stress resistance, and motor activity, displayed in the SOD (superoxide dismutase) mutant flies (31). Another interesting study also indicates the heterogeneity of social environment mediates the cancer progression in the Drosophila model (32). Consistent with their discoveries, the lifespan of fruit flies living in the same space was affected by the compositions of aged and young adults (Fig. 1). In this study, we first demonstrated the existence of “youth impacts” and their capability to promote the increase of lifespan in aged flies. To benefit from the youth impacts, the number of co-housed young flies in the same living space had to be greater than the aged flies (e.g., Fig. 1, ratio 1:3 and 1:9). Presumably, the signals of youth impacts have to be sufficiently large enough to initiate the pro-longevity effects in the aged flies. Furthermore, to perceive the most beneficial effects of youth impacts, there was a best chronological window for aged flies to make such social connections with young flies. As indicated in Figs. 1 and 2, the lifespan extending effect of young impacts was most profound in flies of 40d-old, but not in flies of younger or older ages. These results suggest the sensory systems within the 40d-old flies may be particularly responsive to the youth impacts. Also notably, the youth impacts were expired very soon after 10 days of adult life (Fig. 3). For the aged flies to perceive the full effects of youth impacts, the sensory inputs from olfaction and gustation were essential (Fig. 4). Therefore, the functional integrity of olfactory and gustatory sensing systems in the aged flies may determine the efficacy of youth impacts. Consistently, diverse sensory modalities and associated neural circuits have been implicated in modulating the energy homeostasis and longevity of organisms (5, 6). Based on our results, the most profound effects of longevity promotion by the youth impacts were around 14 to 25% of increase (Fig. 1). Comparing to efficacy of known pro-longevity manipulations (34-36), the overall effects provided by youth impacts were modest. However, similar to the lifespan extension mediated by the reduction of caloric intake (37, 38), the elicitation of youth impacts does not involve the manipulation of endogenous/ectopic genetic element(s) or drug treatment. Instead, the youth impacts work through the variations of social surroundings and the contents of social engagement, representing a more natural pro-longevity approach. Furthermore, multiple lines of evidence suggest the social perception mechanisms are mostly mediated by smell- and taste-dependent chemical cues, such as pheromones and additional volatile/non-volatile chemicals found on the fly body. Intriguingly, earlier studies have demonstrated the chemical compositions of know pheromones vary significantly in fruit flies of distinct chronological ages (6, 28). Therefore, the unique pheromone profiles of young fruit flies may act as an important source of youth impacts and the sensory perception to these chemicals may directly or indirectly influence the physiology and aging process of aged flies. Moreover, it is still not clear if the youth impacts are effective between different Drosophila species, given the pheromone compositions show distinctive features among Drosophila species (39, 40). More interestingly, the human body odors are found to be able to convey a variety of biological and social information to the surrounding individuals. Comparable to the cuticular substances found on the body of fruit flies, the chemical compositions of human body odor are also extremely complicated and change in an age-dependent manner (41, 42). One such example is the increase of 2-Nonenal (C_9_H_16_O), an unsaturated aldehyde with an unpleasant greasy and grassy odor, in the body of elders, suggesting its involvement of the age-related change of body odor (43). Also consistently, a previous study has confirmed the connection between social network size and olfactory function (44, 45). Future studies to identify and characterize chemicals from the scents emanated from young organisms (including the young fruit flies) that possess beneficial/pro-longevity capability will help us uncover the mechanisms of youth impacts and roles of social perception.

It is known that sensory perception has direct influences on the physiology and the aging progression (5, 6, 33). Animals not only passively experience the changes of diverse external and internal states, but also evaluate and react according to changes. The key internal states include the concentrations of metabolites, the energy balance, and the osmolarity of body fluid. On the other hand, the perceptive experiences to social surroundings, time, and potential stresses/threats, provide the links of external environments to the internal physiology and neural processing. Particularly, many clinical studies have also associated the favorable social surroundings with the increase of overall well-being and the reduction of distinct aging-related diseases (10, 46, 47). Here, our results using the Drosophila model system demonstrated the longevity promoting capability of youth impacts derived from young flies of the same gender. Albeit the effects of youth impacts from the opposite gender are not tested in this study, earlier studies indicate the sexual activities and functional fertility could incur a survival cost (6, 48-50). However, intriguingly, successful copulation can endow the male flies with beneficial effects (6), suggesting sex behaviors/experiences may differentially influence the life expectancy of male and female flies.

The definition of healthy aging is to allow the elderly individuals to maintain proper functions in physiology, mental status, and social well-being. Addition to the advances of medical care and disease prevention, in the modern human society, studies by social scientists also suggest cross-generational co-housing, which brings active social engagements between elders and younger adults, is a promising and viable approach to promote healthy aging of elders, highlighting the influences of social surroundings (51, 52). Here, our study in the Drosophila model provided extensive experimental evidence that not only substantiates the notion that social perception/interaction has a great impact on the life expectancy of aged individuals, but also points out potential limitations and requirements for the best efficacy of youth impacts.

## Material and Methods

### Drosophila strains

Flies were reared with standard cornmeal-yeast-agar medium at 25°C, and on a 12/12 hr light/dark cycle. The following Drosophila lines were used: w1118 (BDSC 3605), Canton-S (BDSC 64349), Oregon-R (from Dr. Chun-Hong Chen, National Health Research Institutes, Taiwan), or83b mutant (BDSC 23130; w[*]; TI{w[+m*]=TI}Orco[2]), ppk23 mutant (BDSC 12571; w[1118]P{w[+mGT]=GT1}ppk23[BG01654]).

### Lifespan assay

Unmated flies were collected and separated by gender soon after eclosion. Twenty flies were sorted in a vial and incubated at 25°C. Flies were transferred to a new vial containing fresh medium every 2-3 days. Number of the dead flies was counted and recorded. Survival analyses were performed using the Prism 7 (GraphPad Software). To explore the effects of cuticular extracts, a single fly was cultured in a vial with a filter paper (1 cm^2^) rinsed with isolated cuticular extracts from three 1d-old young flies.

### Co-housing experiments

Unmated flies were collected and sorted into 20 flies per vial soon after eclosion. After 2-3 days, the right wing of flies were clipped to remove the portion beyond the rear end of the abdomen under CO_2_ anesthetization. The marked flies were used as the experimental aged flies. At the indicated dates, aged flies and unmated young flies were cultured in the same vial at the specified ratio (aged:young; 1:9, 1:3, 1:1, 3:1) and keep the population density at 20 flies/vial. Death of co-housed aged and young flies were counted and recorded every 2-3 days. For the controls of the co-housing experiments, the age-matched siblings were recruited to the control vial at the indicated dates to keep the same population density at 20 flies/vial. Deaths of control flies were recorded along with the experimental groups. Statistic analyses were performed with the Prism 7 software (GraphPad Software).

### Collection of cuticular extracts

Fifteen 1d-to 2d-old young flies were collected and soaked into hexane (10 µL per fly; Sigma #32293). After one hour incubation on the rocking shaker at room temperature, hexane was removed by the water circulating vacuum pump. The dried cuticular extracts were stored at -20°C. Right before the experiments, the cuticular extracts were resolved with 30 µl hexane and applied onto a 1cm×1cm filter paper (Whatman #1001-090). The filter paper was then placed into the vial contain fresh fly food. A single fly was introduced into the conditioned vial and transferred to a new vial every 2 days. Also note, during the assay, the living space of the single-housed fly was reduced to half by adjusting the height of cotton stopper. Restriction of the living space was to increase the frequency of physical contact with the filter paper.

### Quantification and statistical analysis

Statistical tests were conducted using Prism 7 (GraphPad Software). The lifespan of aged flies was plotted as scatter plot with mean lifespan ± SEM and the differences were evaluated by Mann-Whitney test. The survival curves were plotted as Kaplan Meyer plots and the statistical significance was tested using the log-rank (Mantel-Cox) test.

## Acknowledgements

We thank Dr. Chun-Hong Chen for the Oregon-R fly line. We also thank the Bloomington Drosophila Stock Center (IN, USA) and Fly Core in Taiwan for providing and handling the fly stocks. This manuscript was aided by comments and discussion from Dr. Yan-Hwa Wu Lee, Dr. Tsai-Wen Chiu, and Dr. Tse-Chun Kuo.

## Funding

This work was supported by the Ministry of Science and Technology, Taiwan (MOST104-2628-B-009-001-MY3 to C.-F. K. and MOST107-2320-B-009-003-MY3 to C.-F. K.) and the “Center For Intelligent Drug Systems and Smart Bio-devices (IDS^2^B)” from The Featured Areas Research Center Program within the framework of the Higher Education Sprout Project by the Ministry of Education (MOE) in Taiwan.

## Author contributions

L.-C. C. and C.-F. K. designed the experiments. L.-C. C. and C.-C. Y. performed all the genetic and behavioral assays, as well as the data analysis. Figures were prepared by L.-C. C. and C.-C. Y. L-C. C and C.-F. K interpreted the results and wrote the manuscript.

## Competing interests

Authors declare no competing interests.

## Data and materials availability

All data are available in the manuscript or the supplementary materials; raw data are available upon request.

## Supplementary Materials

### Supplemental Figures

**Fig. S1.**
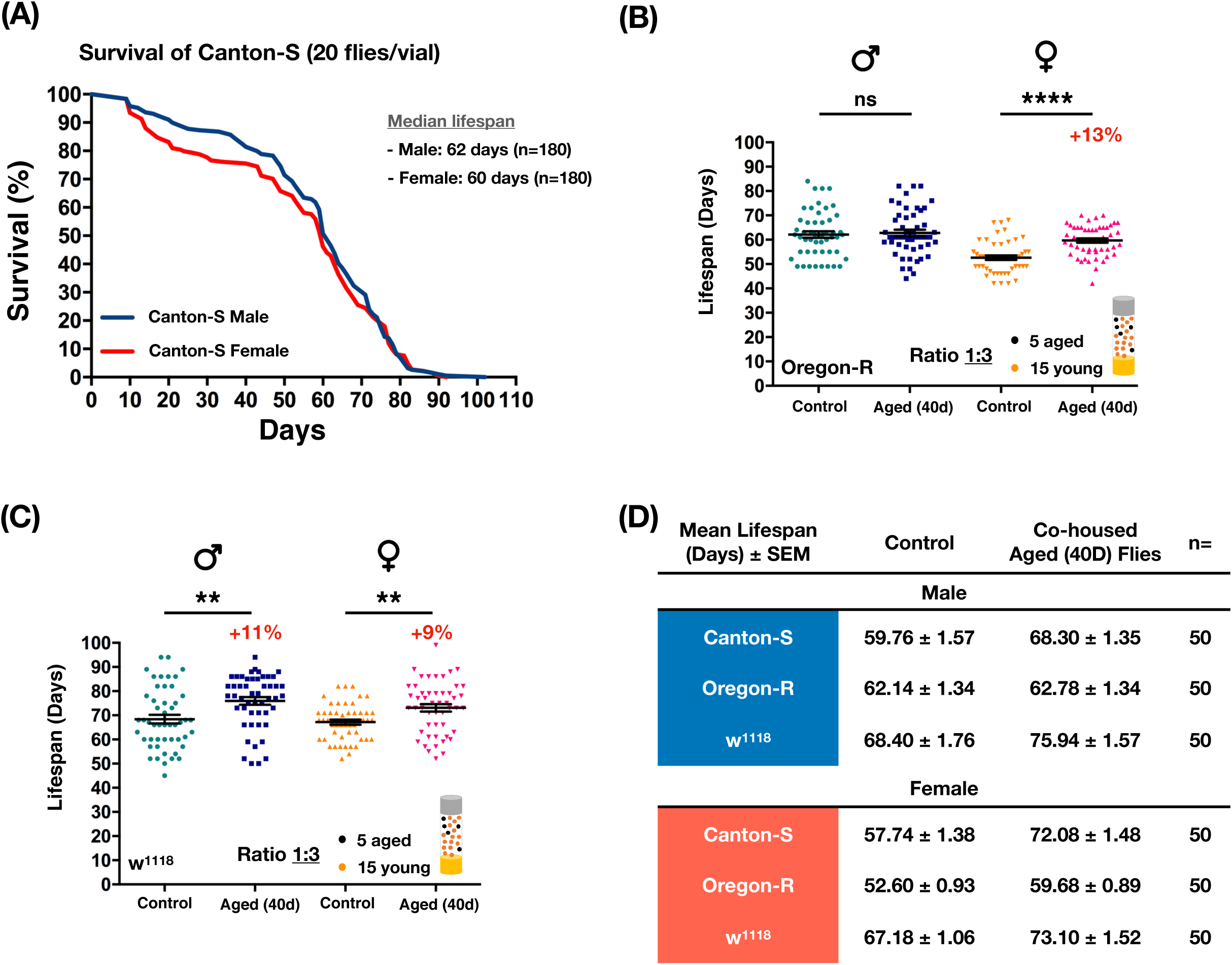
The youth impacts can be observed in w^1118^ and Oregon-R flies. **(A)** Survival curves of the Canton-S flies. Flies were kept at the density of 20 flies per vial and their survival was recorded every two days. The median lifespan of Canton-S was about 62 days in male flies and 60 days in female flies (n=180). **(B)** 40d-old Oregon-R or **(C)** w^1118^ flies were co-cultured with 1d-old conspecific young flies at the ratio of 1:3. The lifespan of individual flies was recorded and analyzed. In both WT lines, the mean lifespan of aged flies was notably increased to different levels, except the case of Oregon-R male flies. Results were expressed as means ± SEM. Please see Fig. 1C for the summary of lifespan results shown in (B) and (C). **p<0.01, ****p<0.0001. p values were annotated by the Mann-Whitney test. Each column contains results of 50 flies. ns: not significant.

**Fig. S2.**
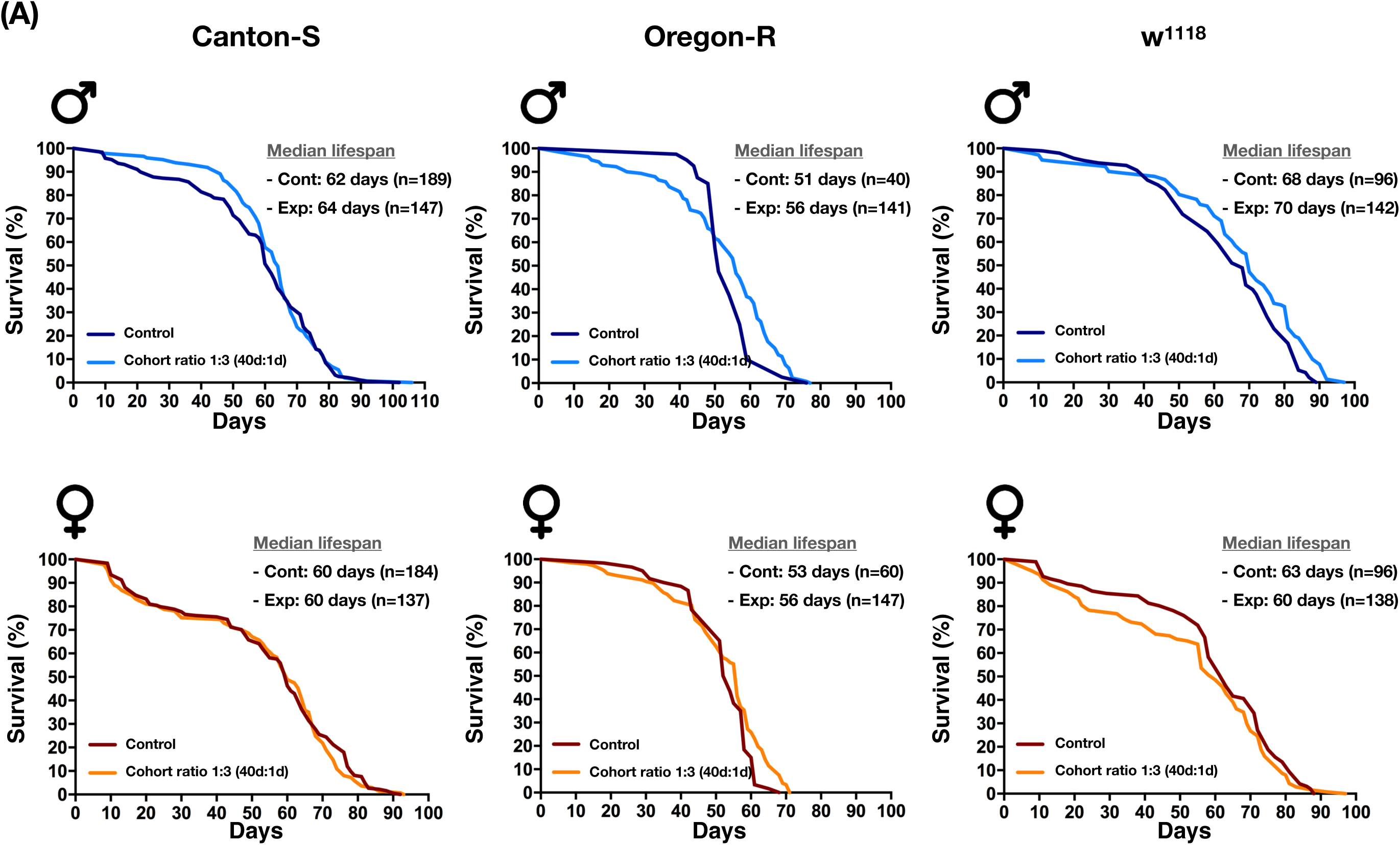
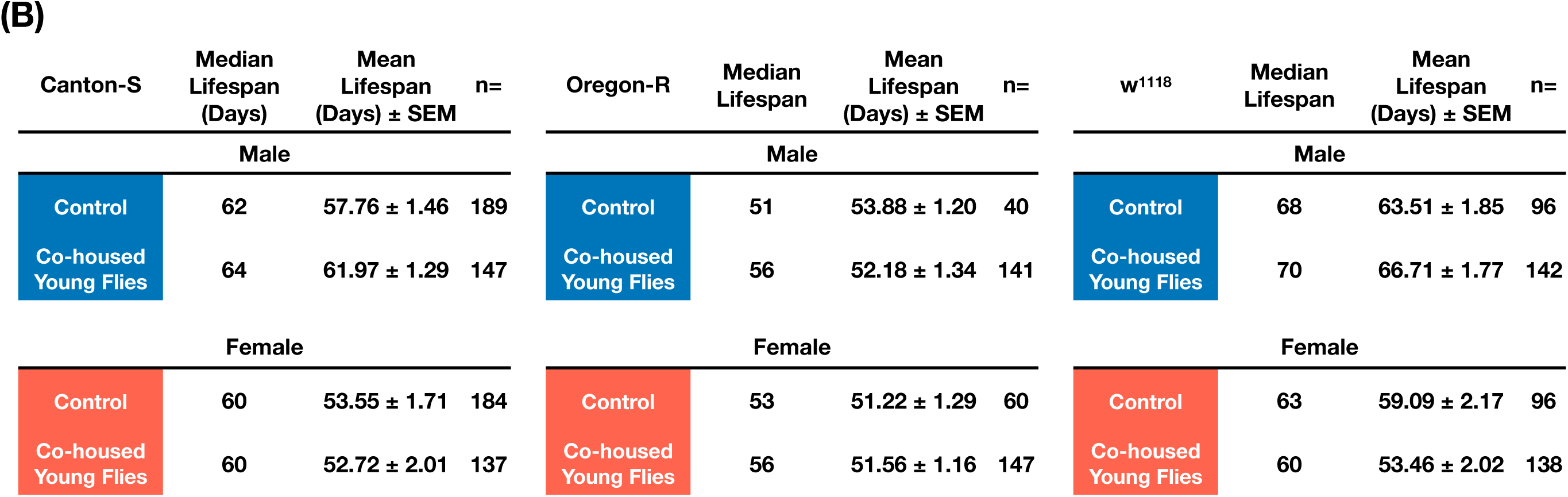
The lifespan of experimental co-housed young flies is not affected by the aged flies. **(A)** Survival curves of the experimental 1d-old young flies (Exp) (three WT species were shown: Canton-S, Oregon-R, and w^1118^) that were co-housed with 40d-old aged flies at the 1:3 ratio had no significant differences compared to the control flies (Cont) that were maintained at a density of 20 flies per vial. p values were annotated by the Log-rank (Mantel-Cox) test. In all cases, p values were greater than 0.05. **(B)** Summary of results shown in (A).

**Fig. S3.**
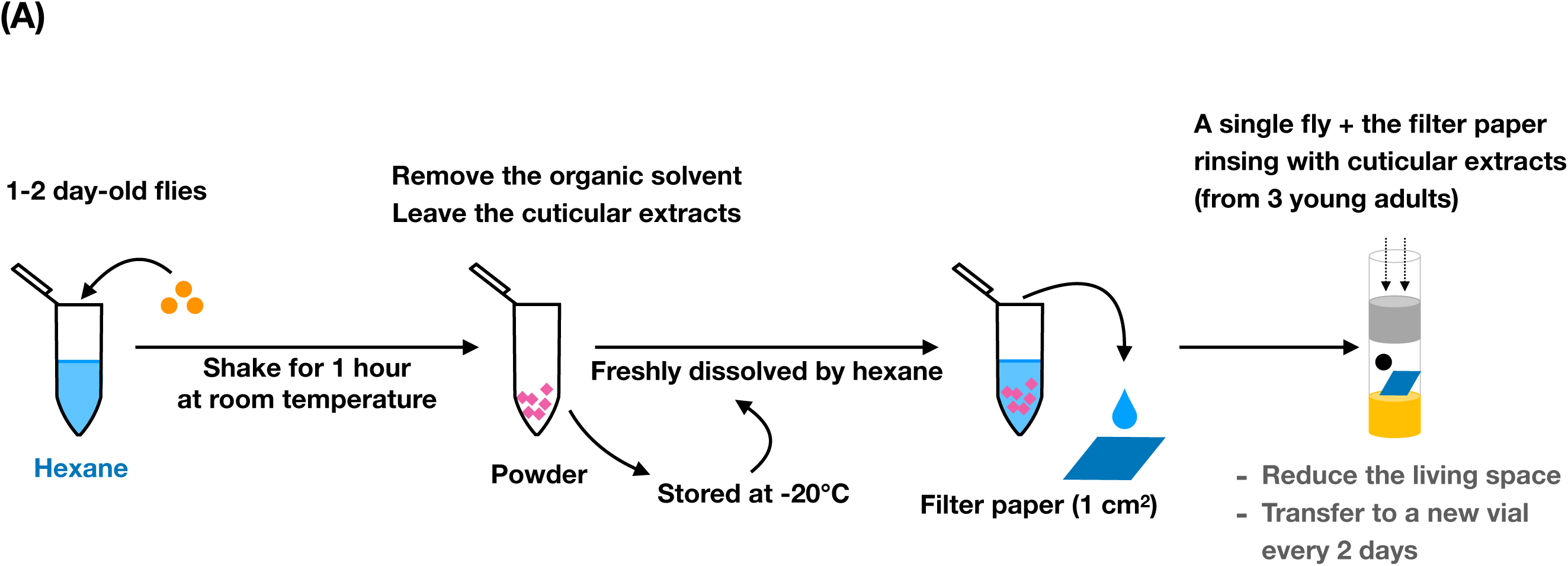
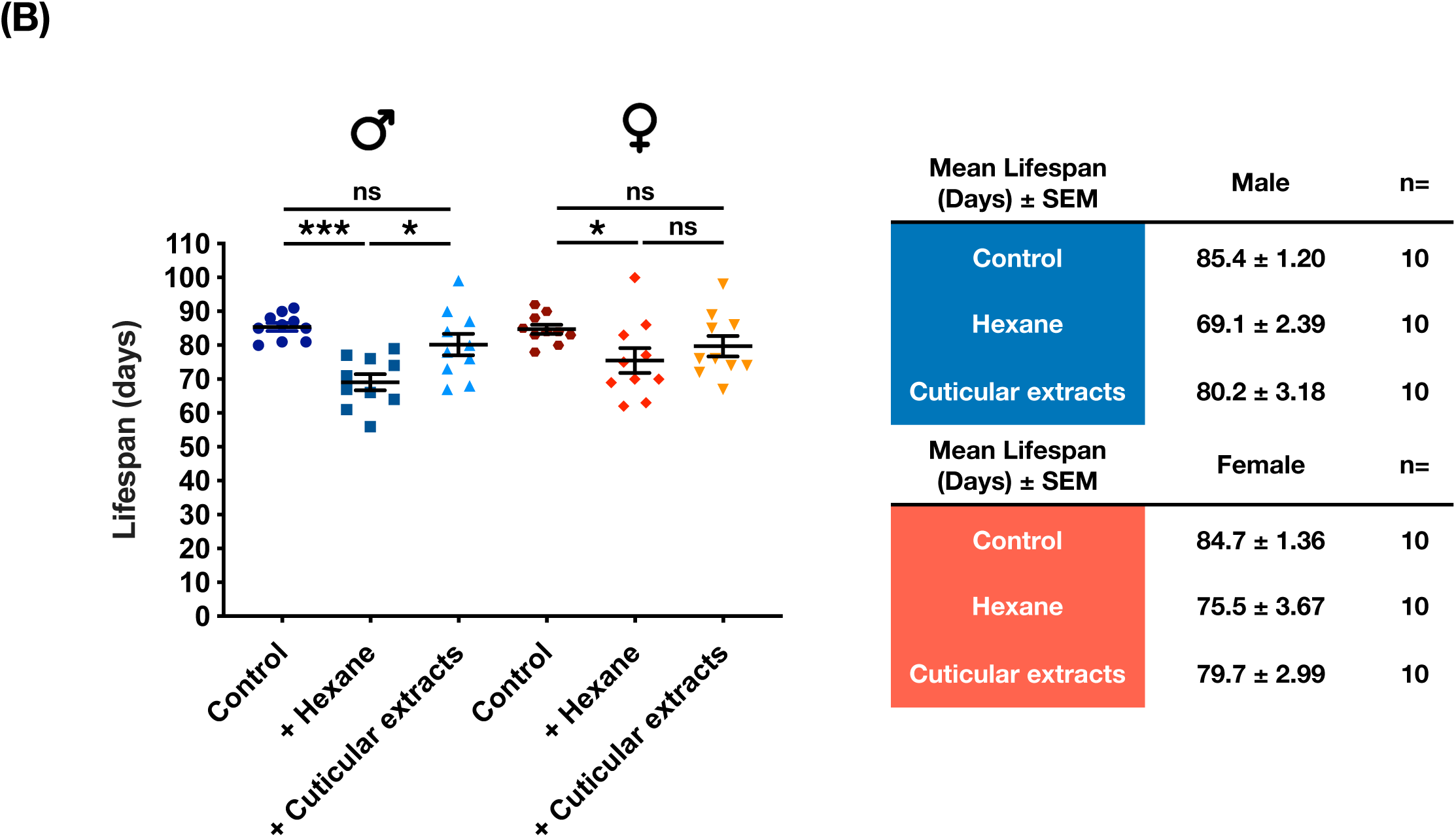
Explore the lifespan modulatory effects of cuticular extracts collected from young flies. **(A)** The experimental illustration describes the processes of retrieving cuticular extracts from 1∼2d-old Canton-S flies and the subsequent culturing conditions. **(B)** To explore the potential source of youth impacts conferred by the young flies, a single Canton-S fly was cultured in a vial containing the regular cornmeal food and a piece of filter paper rinsed with cuticular extracts isolated from three 1d to 2d-old same gender flies (+ Cuticular extracts). The fly was transferred to another vial containing cornmeal food and the freshly prepared cuticular extract-rinsed filter paper every two days. For the hexane control experiments, a single fly was cultured with the filter paper rinsed with hexane (+ Hexane). Another control experiment included only a single Canton-S fly without any treatment (Control). Results were expressed as means ± SEM. *p<0.05, according to Mann-Whitney test (each column, n=10). ns: not significant.

## Notes

### Competing Interest Statement

The authors have declared no competing interest.

